# Decoupling CAR-T Expansion, Conversion, and Decay Timing: Physiologically Aligned Semi-Mechanistic Modeling with Smooth Gating and a Cauchy Likelihood Residual Model

**DOI:** 10.64898/2026.03.01.708827

**Authors:** Yiming Cheng, Yan Li

## Abstract

Chimeric antigen receptor (CAR) T-cell therapies exhibit complex cellular kinetics characterized by rapid expansion, contraction, and long-term persistence, often with substantial inter-individual variability, frequent below-quantification (BLQ) observations, and occasional influential data points. These features can destabilize inference under Gaussian residual assumptions and motivate robust, likelihood-based approaches that can jointly accommodate outliers and censoring. In prior work, we showed that combining Student’s *t* residuals with likelihood-based BLQ handling (M3 censoring) improves robustness in CAR-T cellular kinetics modeling; however, implementing Student’s *t* censoring likelihoods is not straightforward in all modeling platforms because the Student’s *t* cumulative distribution function (CDF) lacks a simple elementary closed-form.

Here, we evaluated the Cauchy residual likelihood as a practical, heavy-tailed alternative to Student’s *t* that provides closed-form expressions for both the probability density function (PDF) and CDF. In a two-compartment IV pharmacokinetic simulation with terminal-phase outlier contamination, Cauchy residuals preserved stable parameter recovery comparable to Student’s *t* while avoiding the bias and instability observed under Normal residuals. In a real-data integrated CAR-T cellular kinetics application fitted with full Bayesian inference and likelihood-based BLQ handling, replacing Student’s *t* with Cauchy yielded highly concordant posterior inference and similar subject-level predictions.

We further extended a semi-mechanistic CAR-T cellular kinetics framework by replacing piecewise switching with smooth S-shaped time-varying rate functions and allowing process-specific transition times. Full Bayesian posterior summaries supported asynchronous transition timing, including earlier conversion relative to expansion and later decay-related transitions. Collectively, these results support Cauchy likelihoods as an implementation-friendly robust option and demonstrate that smooth, process-specific transition modeling can enhance physiological plausibility for CAR-T kinetics.

## Introduction

Chimeric antigen receptor (CAR) T-cell therapies have reshaped the treatment of several hematologic malignancies by enabling engineered T cells to expand, contract, and persist in vivo after a single infusion. ^1-6^ Unlike small molecules and monoclonal antibodies, CAR-T products behave as “living drugs,” and their cellular kinetics reflect a combination of proliferation, phenotypic evolution, and cell loss rather than classical absorption and clearance. ^7-12^ This multi-phase behavior—typically rapid expansion followed by contraction and long persistence—shows substantial inter-individual variability and creates practical challenges for quantitative modeling. ^9^ Robust population cellular kinetics models are therefore needed to support interpretation of exposure–response relationships, characterize variability, and improve the reliability of inference across trials and products. ^13^

Two recurring issues limit the stability and portability of current CAR-T modeling workflows. First, CAR-T datasets often contain influential observations and extreme deviations, particularly during early expansion and late persistence, ^8-11, 14, 15^ that can destabilize parameter estimation under Gaussian residual assumptions. Second, transgene measurements are typically generated by qPCR assays spanning multiple orders of magnitude, ^16^ resulting in a substantial fraction of observations below the lower limit of quantification (BLQ), especially during contraction and persistence. ^10, 11, 15^ Because BLQ values still contain information, they should be incorporated through likelihood-based censoring rather than omitted or imputed. ^17-23^ In practice, CAR-T modeling frequently requires methods that can handle outliers and censoring simultaneously. ^9, 13^

Our prior work showed that integrating Student’s *t* residuals with likelihood-based BLQ handling (M3 censoring) in a full Bayesian framework improves robustness and uncertainty quantification. ^9, 13^ However, Student’s *t* likelihoods can be challenging to deploy across platforms when censoring contributions require CDF evaluation, particularly in environments such as Monolix that do not natively support the Student’s *t* CDF in user-defined likelihood implementations. This creates a practical barrier to cross-software reproducibility and motivates evaluation of heavy-tailed alternatives that preserve robustness while simplifying implementation.

In this study, we evaluate the **Cauchy** residual likelihood as a practical alternative to Student’s *t*. Cauchy is a special case of the Student’s *t* family (ν = 1) and has simple closed-form expressions for both the PDF and CDF, which materially reduces implementation burden for robust likelihoods and censoring components. ^24, 25^ The central methodological question is whether Cauchy can reproduce the robustness behavior of Student’s *t*—without introducing distortion under clean data—while improving portability across modeling platforms.

A second focus of this work is model structure. Widely used semi-mechanistic CAR-T cellular kinetics models commonly rely on piecewise step functions with a shared transition time to represent switching between expansion, conversion, and decay processes. ^5, 7, 9-12, 15^ While convenient, this structure assumes instantaneous and synchronized transitions, which may be physiologically unrealistic given that T-cell proliferation, differentiation, and loss occur gradually and may be temporally decoupled. To address this limitation, we propose a smooth-gating framework that replaces discontinuous switching with S-shaped time-varying rate functions and allows process-specific transition times, enabling asynchronous timing across expansion, conversion, and decay.

The objectives of this study were: (1) to compare Cauchy, Student’s *t*, and Normal residual likelihoods under controlled terminal-phase outlier contamination in a two-compartment IV PK simulation; (2) to test whether Cauchy reproduces Student’s *t* inference in an integrated CAR-T kinetics application under full Bayesian estimation with likelihood-based BLQ handling; and (3) to evaluate a smooth-gating, process-specific transition CAR-T model and quantify timing separation among expansion, conversion, and decay using posterior uncertainty. Together, these contributions aim to strengthen robustness, improve physiological plausibility, and enable more practical cross-platform implementation for CAR-T cellular kinetics modeling.

## Methods

### Modeling platforms and cross-software implementation

All data assembly and post-processing were performed in RStudio (R v4.1.3; Posit Software, Boston, MA, USA). Model fitting was conducted in **Monolix®** (2023R1; Lixoft SAS, Antony, France) and **NONMEM®** (7.5.0; ICON Development Solutions, North Wales, PA, USA). NONMEM was specifically used for **full Bayesian confirmation** via MCMC to characterize posterior uncertainty and evaluate robustness of inference under alternative likelihood specifications. Uninformative priors were used throughout: fixed effects were assigned multivariate normal priors with large variances, and covariance structures for random effects were assigned inverse Wishart priors. For standard continuous outcomes, Monolix does not natively support user-defined residual likelihoods in the same way as NONMEM. To enable custom residual-error likelihood evaluation in Monolix, we implemented a workaround using the count-model interface while preserving a continuous-data likelihood. Specifically, observed concentrations were **scaled by 10**□ **and rounded to integers** prior to import so that Monolix would accept the dependent variable as a count outcome. Within the model code, the integer observations were immediately mapped back to the original continuous scale by dividing by **10**□, and the intended continuous residual likelihood (e.g., Normal, Student’s *t*, Cauchy) was evaluated on the rescaled values. The **10**□ factor was selected to retain numerical precision during integer encoding while maintaining stable likelihood evaluation and ensured rounding error was negligible relative to assay variability.

A practical motivation for using Monolix and adopting a cross-software workflow was to reflect a realistic implementation constraint: while NONMEM readily accommodates many flexible likelihood functions for pharmacometricians, deploying robust residual models in Monolix typically requires additional engineering. In this context, the Cauchy likelihood—with closed-form PDF and CDF expressions—was evaluated as a portability-focused alternative that can support robust inference (including censored contributions when needed) with minimal implementation overhead across modeling environments.

### Data Simulation

To compare the performance of a Cauchy residual likelihood with a Student’s t likelihood for robust handling of outliers (and, where relevant, censored contributions), we conducted a dedicated simulation study in which datasets were generated under a standard two-compartment IV PK model and then refit using alternative residual-error distributions. IV bolus two-compartment PK profiles were simulated for 25 subjects using typical parameter values: CL = 2.3 L/h, V1 = 20 L, Q = 5 L/h, and V2 = 70 L. Inter-individual variability (IIV) was assumed log-normal with 20% variability applied to all structural parameters, and proportional residual error was assumed with σ = 20%. No covariate effects or additional random effects were included. Each subject received a single 5000 mg IV bolus dose. Concentrations were sampled at 0, 0.125, 0.25, 0.5, 1, 2, 4, 6, 8, 10, 12, 14, 16, 18, 20, 24, 36, and 48 h, providing adequate characterization of distribution and the terminal phase under the clean-data scenario.

To introduce controlled outlier contamination while preserving the underlying PK trajectory, we perturbed a single terminal-phase observation (48 h) by multiplying the simulated observation by a prespecified factor. A factor of 1 represented the clean baseline scenario. Factors of 6 and 10 were used to represent moderate deviations, and 15 and 20 to represent extreme deviations, designed to mimic influential terminal observations capable of biasing clearance-related inference. Each simulated dataset was refit under three residual-error likelihoods: Normal (Gaussian), Student’s t, and Cauchy. Performance was compared across contamination severities based on recovery of population and individual parameters, supplemented by visual assessment of representative individual concentration–time profiles with observations overlaid on model predictions.

### Clinical Study Data

This analysis used data from three CAR-T clinical trials—TRANSCEND14, KarMMa-3, and EVOLVE—as summarized in our previous publication. ^10, 11, 15^ Across these studies, baseline demographic and disease characteristics were broadly similar, suggesting generally comparable enrolled populations.

All studies were conducted in accordance with the Declaration of Helsinki, International Council for Harmonisation (ICH) Good Clinical Practice (GCP) guidelines, and applicable local regulatory requirements. Study protocols and any subsequent amendments were reviewed and approved by the relevant institutional review boards or independent ethics committees at participating sites, and all participants provided written informed consent prior to study procedures.

### Residual-error likelihoods evaluated

Three residual-error probability density functions (PDFs) were evaluated. The Normal (Gaussian) model (Eq. 1) corresponds to a squared-error loss and therefore penalizes large deviations quadratically. The Student’s *t* model (Eq. 2) estimates the degrees of freedom, *v*, and exhibits power-law tail decay, providing increasing robustness to extreme deviations as *v* decreases. The Cauchy model (Eq. 3 for the PDF and Eq. 4 for the CDF) is a special case of the Student’s *t* distribution with *v* = 1. Notably, both the Cauchy PDF and CDF have simple closed-form expressions, enabling straightforward implementation in Monolix. In contrast, the Student’s *t* CDF does not have a closed-form expression, which precludes direct implementation of the corresponding CDF-based likelihood in Monolix.

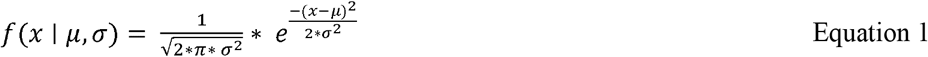

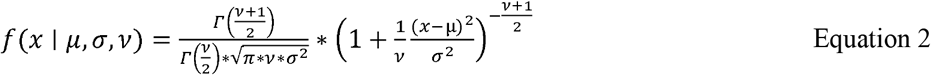

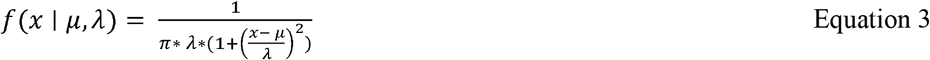

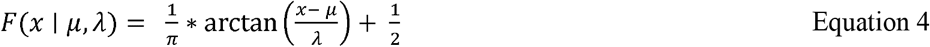

Here, *x* denotes the observed value and *μ* denotes the model-predicted value. *σ* is the standard deviation for the Normal and Student’s *t* likelihoods, *λ* is the scale parameter for the Cauchy likelihood, and *v* is the degrees of freedom for the Student’s *t* likelihood. Γ(·) denotes the gamma function and arctan(·) denotes the arctangent function.

### Population cellular kinetic structure model

Three semi-mechanistic population cellular kinetic (CK) structures were evaluated side-by-side. The reference model was the widely used piecewise shared-transition framework derived from the theoretical work of De Boer and Perelson on vigorous immune responses (Figure 1). ^26^ In this construct, CAR-T kinetics are represented by an initial expansion phase followed by a biphasic contraction phase. During early expansion, activated/effector-like CAR-T cells proliferate rapidly under a non–antigen-limited growth assumption for a finite period. After the transition, the system enters contraction, during which activated CAR-T cells either die rapidly or convert into longer-lived memory-like CAR-T cells. The activated and memory-like compartments then decline with their own decay rate constants.

**Figure 1.**
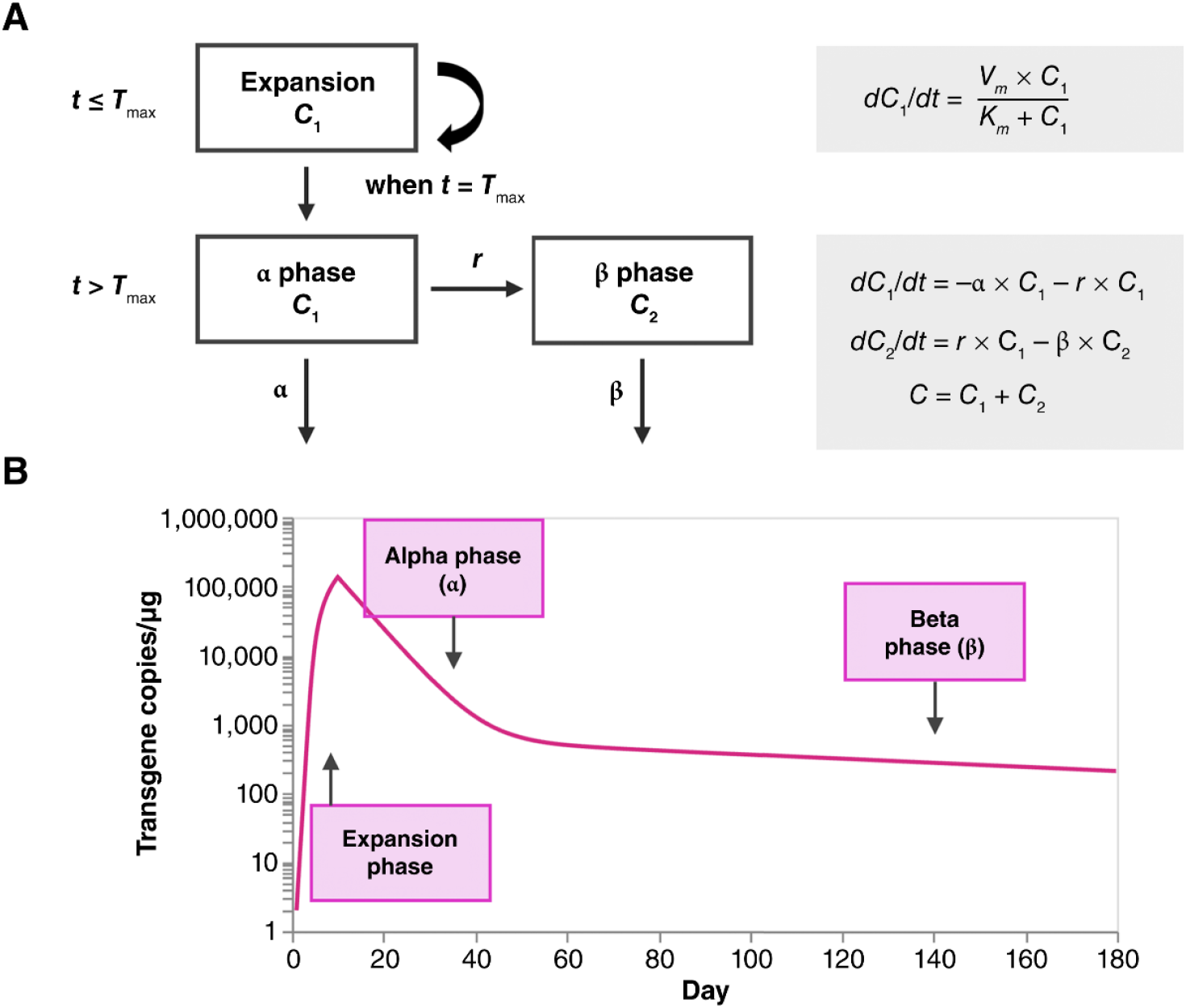
Semi-mechanistic cellular kinetics model of CAR-T therapies. (**A)** Compartmental model and the associated differential equations that describe T-cell kinetics during the expansion and contraction phases. (**B)** Graphical representation of the typical cellular kinetics profile. *C*_1_, transgene levels in expansion phase (time < *T*_max_) or α contraction phase (time ≥ *T*_max_); *C*_2_, transgene levels in β contraction phase; *Km*, transgene levels giving expansion rate of 1/2 *Vm*; *r*, conversion rate constant from rapid contraction phase to slow contraction phase; *T*_max_, time to maximum transgene level; *Vm*, maximum expansion rate; α, rate constant of rapid contraction phase; β, rate constant of slow contraction phase.

In our prior work, we modified this traditional semi-mechanistic model by introducing saturable expansion, in which pre-transition proliferation is governed by a *v*_max_/*κ*_*m*_-type formulation; this modification improved model fit, particularly at early time points. ^11, 15^ In the current work, we further extended the framework by converting the piecewise transition structure into a smooth-gating, process-specific transition model. This formulation replaces discontinuous piecewise switching with continuous, S-shaped time-varying rate functions and relaxes the constraint that distinct biological processes share a single synchronized transition time.^27^ This extension was motivated by the expectation that expansion, conversion, and decay processes need not change instantaneously or simultaneously.

In the smooth-gating specification, time-varying rate functions were defined for activated-cell expansion *rho* (*t*), conversion to memory-like cells *r* (*t*), activated-cell decay *α* (*t*), and memory-like cell decay *β* (*t*) using Hill-type S-shaped transitions (Equations 5–8):

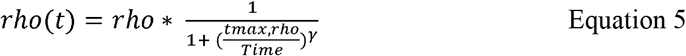

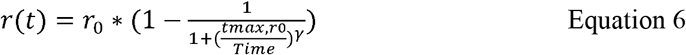

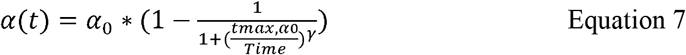

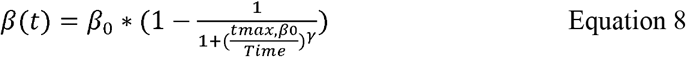

Here, rho denotes the maximal activated T-cell expansion rate and *t*_max_ ,_*rho*_ is the characteristic time at which expansion transitions toward cessation. *r*0denotes the maximal conversion rate from activated to long-lived memory-like T cells and *t*_max_ ,_*r*_ is the characteristic time at which conversion turns on. *α*_o_ denotes the maximal activated T-cell decay rate and *t*_max ,*α*_is the characteristic time at which activated-cell decay turns on. *β*_0_ denotes the maximal memory-like T-cell decay rate and *t*_max ,*β*_ is the characteristic time at which memory-like cell decay turns on. The Hill coefficient *gamma* (γ) controls the steepness of each transition.

## Results

### Simulation case study (Monolix SAEM): Cauchy Residuals Provide Outlier Robustness Comparable to Student’s t

Previously, we demonstrated that a Student’s *t* residual model can improve the accuracy and precision of population- and individual-level parameter estimates in the presence of outlier contamination. ^9, 13^ Here, we performed an analogous simulation to evaluate whether a **Cauchy** residual likelihood can deliver comparable outlier robustness while offering closed-form PDF and CDF expressions that simplify implementation across modeling platforms. Briefly, data were simulated under a two-compartment IV PK model, and population parameter estimation was performed under three residual likelihoods: **Normal, Student’s *t***, and **Cauchy**.

Outlier/contamination severity was controlled using a multiplicative-factor scheme, in which a single terminal-phase observation at 48 h post-dose was multiplied by a predefined factor (1 = no contamination; 6 and 10 = moderate contamination; 15 and 20 = severe contamination).

Figure 2a summarizes the resulting population parameter estimates (dotted lines indicate the true values). Under clean data (multiplicative factor = 1), the Normal, Student’s *t*, and Cauchy likelihoods produced similar central estimates with overlapping uncertainty for the all PK parameters, indicating that the heavy-tailed likelihoods did not introduce discernible distortion when the data were not contaminated. As contamination increased, the Normal likelihood degraded rapidly, with marked instability and bias in parameters most sensitive to the terminal phase. In particular, CL_pop exhibited pronounced sensitivity (including collapse toward implausibly low values in the most contaminated settings), while Q_pop and V2_pop drifted away from the simulated truth with widening uncertainty. In contrast, the Cauchy likelihood remained stable across the full range of outlier severity, maintaining parameter estimates close to the true values and exhibiting behavior that closely matched the Student’s *t* benchmark under the same settings.

**Figure 2.**
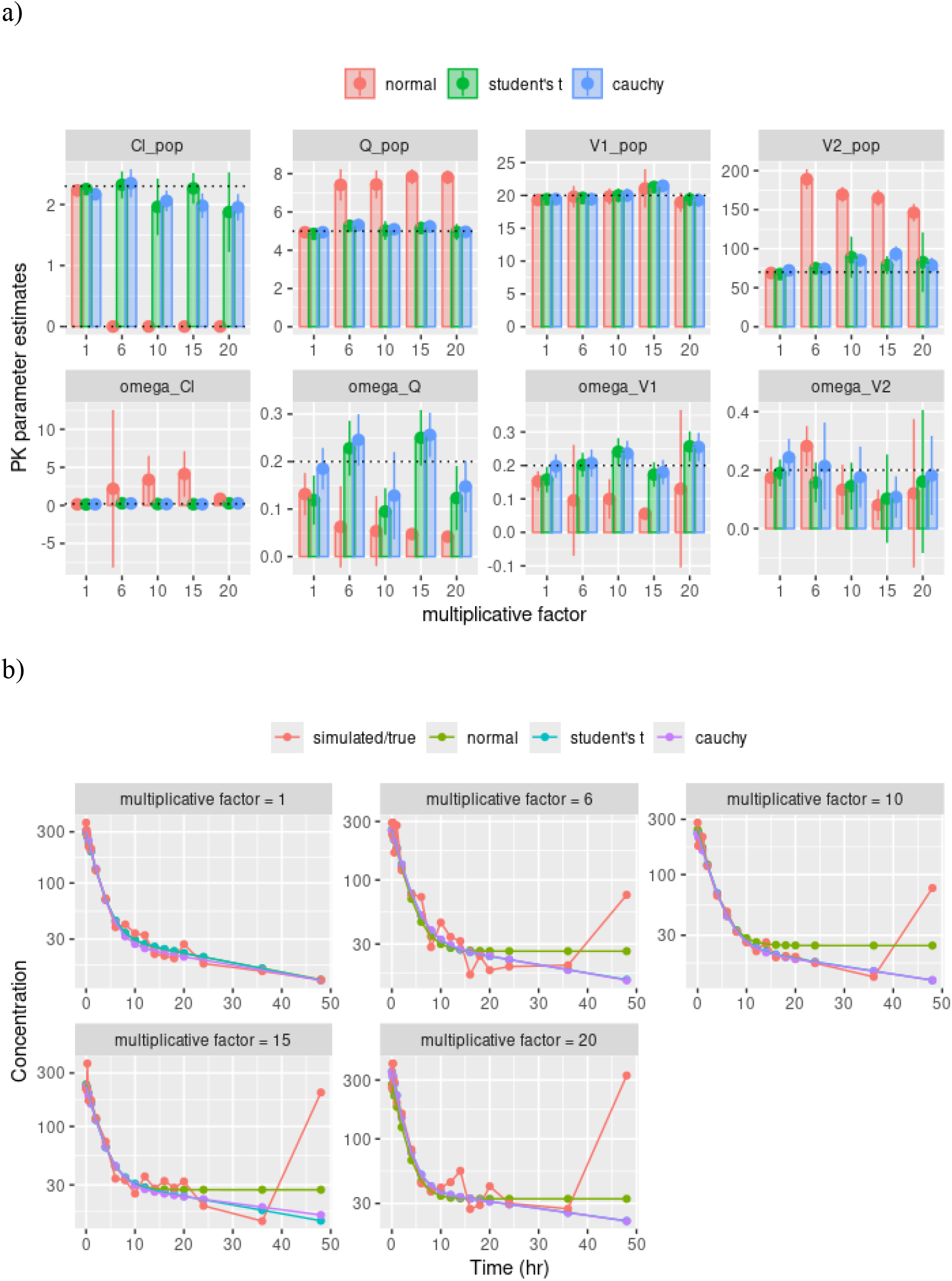
Comparison of Normal, Student’s *t*, and Cauchy residual likelihoods for handling outliers. (a) Population parameter recovery across increasing outlier severity; dotted lines indicate the true simulation values. (b) Individual concentration–time profiles under increasing contamination; dotted lines indicate model predictions. **CL_pop**, clearance; **Q_pop**, inter-compartmental clearance; **V1_pop**, central compartment volume; **V2_pop**, peripheral compartment volume.

Figure 2b provides an illustrative concentration–time comparison for a representative individual under increasing terminal outlier severity. Consistent with the population-level results, the Normal-based fit was visibly distorted by terminal outliers, bending upward to accommodate the contaminated observation, that is, increasing the likelihood assigned to the outlier data point. In contrast, both the Cauchy- and Student’s *t*–based fits remained aligned with the simulated “true” profile across contamination levels, reflecting reduced sensitivity to the high-leverage terminal observation.

Collectively, these simulations show that heavy-tailed residual likelihoods reduce sensitivity to severe terminal outliers relative to Normal residual assumptions in a standard two-compartment IV PK model. Within this simulation design, Cauchy residuals reproduced the robustness behavior of the Student’s *t* benchmark at both the population level (parameter recovery) and the individual level (profile reconstruction).

### Real-data case study (NONMEM full Bayesian): Cauchy versus Student’s t in integrated CAR-T kinetics modeling

To assess whether the simulation findings translate to a realistic complexity setting, we conducted a real-data case study using an integrated CAR-T cellular kinetics model previously developed with a Student’s *t* residual likelihood together with likelihood-based handling of BLQ observations (M3 censoring). ^9^ The objective of this analysis was not to modify the structural model, but to isolate the effect of the residual likelihood by replacing Student’s *t* with a Cauchy likelihood while holding the remainder of the modeling framework constant. Specifically, we retained the same semi-mechanistic structure, including saturable expansion (Vmax/Km), and applied the same BLQ handling approach. Both models were fit using full Bayesian inference in NONMEM to enable a like-for-like comparison of posterior inference.

Figure 3a compares posterior distributions from the Student’s *t* and Cauchy analyses for key fixed- and random-effect parameters. Across the majority of parameters, the posterior distributions show strong agreement between the two likelihoods, with similar locations and substantial overlap of posterior mass. This pattern indicates that, for this integrated CAR-T dataset, replacing Student’s *t* with Cauchy yields comparable posterior inference for the main structural parameters governing timing, expansion, and contraction/persistence dynamics, as well as for most variability components. For a limited subset of parameters, the Cauchy posterior appears modestly more concentrated than the Student’s *t* posterior; however, we interpret this conservatively as a difference in posterior concentration rather than evidence of systematically improved information recovery, because posterior width can be influenced by tail behavior and parameter–error trade-offs. For selected variability terms, posterior modes show small shifts between likelihoods, but these shifts are not accompanied by clear separation at the level of the full distributions shown in Figure 3a.

**Figure 3.**
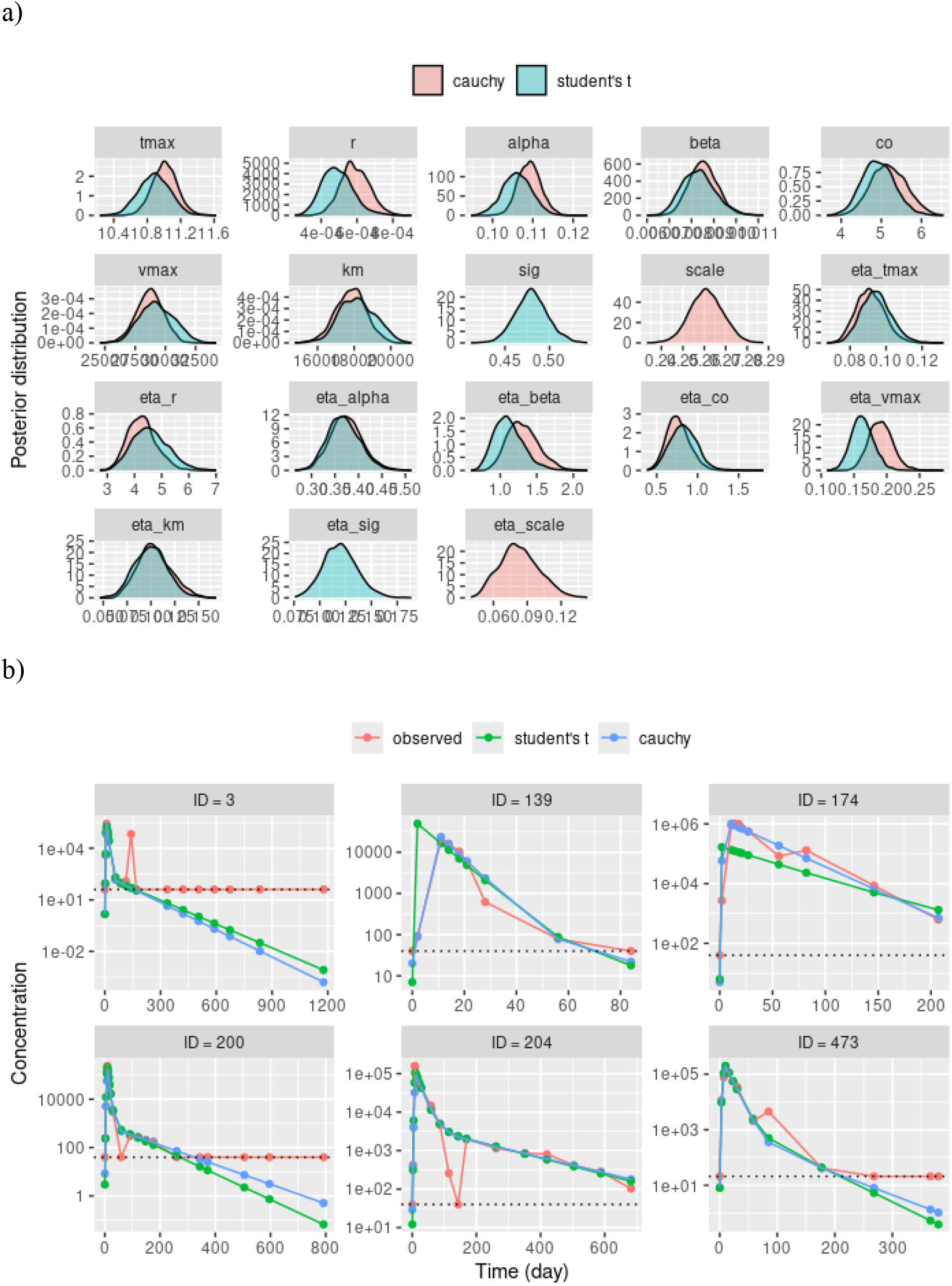
Comparison of posterior inference and predictive performance under Cauchy and Student’s *t* likelihoods for integrated CAR-T kinetics. (a) Posterior distributions of population fixed-effect and variability parameters. **alpha**, rate constant for the rapid contraction phase; **beta**, rate constant for the slow contraction phase; **conc0**, initial transgene level; **km**, transgene level associated with the half-maximal expansion rate (vmax/2); **omega**, inter-individual variability; **r**, conversion rate constant from the rapid contraction phase to the slow contraction phase; **sig**, proportional residual error; **tmax**, time at which T-cell expansion ends; **vmax**, maximum expansion rate; **scale (**λ), scale parameter of the Cauchy distribution. (b) Representative individual concentration–time profiles illustrating predictive alignment; dotted horizontal lines indicate the below limit of quantification (BLQ).

Figure 3b provides representative individual concentration–time profiles to assess whether similarity in posterior inference translates to comparable predictive behavior at the subject level, including regions influenced by BLQ censoring and influential observations. Across the displayed individuals, Cauchy- and Student’s *t*–based predictions are generally similar and track the observed profiles closely, without an evident systematic difference in predictive alignment across subjects. In isolated individuals, one likelihood may show slightly improved local agreement at specific timepoints; however, such subject-level differences are expected in complex nonlinear models and should not be interpreted as a consistent advantage without aggregate predictive diagnostics.

Overall, Figures 3a and 3b indicate that, in this integrated CAR-T cellular kinetics application fitted under full Bayesian inference with likelihood-based BLQ handling, the Cauchy likelihood provides posterior inference and subject-level predictions that are closely aligned with those obtained under a Student’s *t* residual likelihood.

### Smooth time-varying rate constants and process-specific transition times in semi-mechanistic CAR-T cellular kinetics

We extended the commonly used semi-mechanistic CAR-T cellular kinetics framework in two ways: (i) replacing discontinuous piecewise switching with smooth (S-shaped) time-varying rate functions, and (ii) relaxing the constraint that multiple biological processes share a single common transition time (tmax). For this evaluation, models were implemented in Monolix using a Cauchy residual likelihood, and three specifications were compared: (1) a traditional piecewise shared-transition model, (2) a piecewise model with saturable expansion (Vmax/Km), and (3) the proposed model with smooth time-varying rate functions and process-specific transition timing.

In the reference piecewise shared-transition model, a single transition time parameter (tmax1_pop) governs switching behavior across multiple kinetic components. As shown in Table 1, tmax1_pop was estimated at 6.62 days with a narrow uncertainty interval (P2.5–P97.5: 6.46– 6.78 days). Other fixed effects (e.g., rho_pop, alpha_pop, beta_pop) and the proportional residual error term (sig_pop) were estimated with relatively tight uncertainty, and most IIV terms were supported with moderate precision.

**Table 1.** Summary of population cellular kinetic parameter estimates from the traditional piecewise transition model.

| Parameter | Estimate | SE | RSE (%) | 2.5% Percentile | 97.5% Percentile |
| --- | --- | --- | --- | --- | --- |
| tmax1_pop | 6.62 | 0.08 | 1.22 | 6.46 | 6.78 |
| r_pop | 0.00029 | 0.00004 | 13.89 | 0.00022 | 0.00038 |
| alpha_pop | 0.10 | 0.00 | 2.93 | 0.09 | 0.11 |
| beta_pop | 0.0065 | 0.0004 | 6.85 | 0.0057 | 0.0075 |
| conc0_pop | 8.60 | 0.38 | 4.40 | 7.89 | 9.37 |
| sig_pop | 0.27 | 0.01 | 2.43 | 0.26 | 0.28 |
| rho_pop | 1.33 | 0.02 | 1.24 | 1.30 | 1.37 |
| omega_tmax1 | 0.26 | 0.01 | 3.40 | 0.24 | 0.28 |
| omega_r | 2.41 | 0.12 | 5.03 | 2.18 | 2.66 |
| omega_alpha | 0.60 | 0.02 | 4.16 | 0.55 | 0.65 |
| omega_beta | 1.00 | 0.07 | 6.66 | 0.88 | 1.14 |
| omega_conc0 | 0.81 | 0.04 | 4.46 | 0.74 | 0.89 |
| omega_sig | 0.23 | 0.03 | 12.74 | 0.18 | 0.30 |
| omega_rho | 0.25 | 0.01 | 3.55 | 0.24 | 0.27 |

Abbreviations: alpha\_pop, rate constant for the rapid contraction phase; beta\_pop, rate constant for the slow contraction phase; conc0\_pop, initial transgene level; omega, inter-individual variability; r\_pop, conversion rate constant from the rapid contraction phase to the slow contraction phase; rho\_pop, expansion rate constant; sig\_pop, proportional residual error; tmax1\_pop, time at which T-cell expansion ends.

Previously, we showed that replacing a constant pre-transition expansion rate with a saturable Vmax/Km formulation is more physiologically plausible for T-cell expansion and provides improved model fit, particularly at early time points. In the Vmax/Km model (Table 2), ^11, 15^ tmax1_pop increased to 8.65 (8.35–8.96) days. The saturable expansion parameters (vmax_pop and km_pop) were estimated with high precision, as reflected by low relative standard errors (<5%) and narrow uncertainty intervals; the associated variability terms were also supported.

**Table 2.** Summary of population cellular kinetic parameter estimates from the piecewise model with saturable expansion (Vmax/Km).

| Parameter | Estimate | SE | RSE (%) | 2.5% Percentile | 97.5% Percentile |
| --- | --- | --- | --- | --- | --- |
| tmax1_pop | 8.65 | 0.16 | 1.81 | 8.35 | 8.96 |
| r_pop | 0.00049 | 0.00006 | 13.21 | 0.00038 | 0.00063 |
| alpha_pop | 0.12 | 0.00 | 2.89 | 0.11 | 0.12 |
| beta_pop | 0.0077 | 0.0005 | 6.85 | 0.0067 | 0.0088 |
| conc0_pop | 4.20 | 0.28 | 6.67 | 3.69 | 4.79 |
| sig_pop | 0.25 | 0.01 | 2.45 | 0.24 | 0.26 |
| vmax_pop | 35542.95 | 672.87 | 1.89 | 34248.70 | 36886.20 |
| km_pop | 21138.67 | 333.57 | 1.58 | 20495.00 | 21802.50 |
| omega_tmax1 | 0.39 | 0.01 | 3.66 | 0.36 | 0.42 |
| omega_r | 2.33 | 0.12 | 5.03 | 2.11 | 2.57 |
| omega_alpha | 0.59 | 0.02 | 3.65 | 0.55 | 0.63 |
| omega_beta | 1.07 | 0.06 | 5.79 | 0.95 | 1.19 |
| omega_conc0 | 1.01 | 0.06 | 5.45 | 0.91 | 1.13 |
| omega_sig | 0.14 | 0.03 | 21.82 | 0.09 | 0.21 |
| omega_vmax | 0.36 | 0.01 | 3.73 | 0.33 | 0.39 |
| omega_km | 0.22 | 0.01 | 5.89 | 0.20 | 0.25 |

Abbreviations: alpha\_pop, rate constant for the rapid contraction phase; beta\_pop, rate constant for the slow contraction phase; conc0\_pop, initial transgene level; km\_pop, transgene level associated with half-maximal expansion rate (vmax\_pop/2); omega, inter-individual variability; r\_pop, conversion rate constant from the rapid contraction phase to the slow contraction phase; sig\_pop, proportional residual error; tmax1\_pop, time at which T-cell expansion ends; vmax\_pop, maximum expansion rate.

The proposed new model (Table 3) replaces discontinuous switching with smooth S-shaped time variation and allows process-specific transition times, parameterized as offsets relative to the self-expansion transition time. The baseline expansion transition time was estimated as tmax1_pop = 5.57 (5.48–5.67) days. The alpha-decay transition offset (deltmax3_pop) was estimated at 3.06 (2.69–3.44) days, and the beta-decay transition offset (deltmax4_pop) at 9.07 (8.64–9.50) days; in both cases, the uncertainty intervals exclude 0, indicating that the alpha- and beta-decay transitions occurred substantially later than the end of T-cell expansion. In contrast, the conversion transition offset (deltmax2_pop) was estimated at −0.30 day, suggesting a potentially earlier transition from the rapid to the slow contraction phase.

**Table 3.**
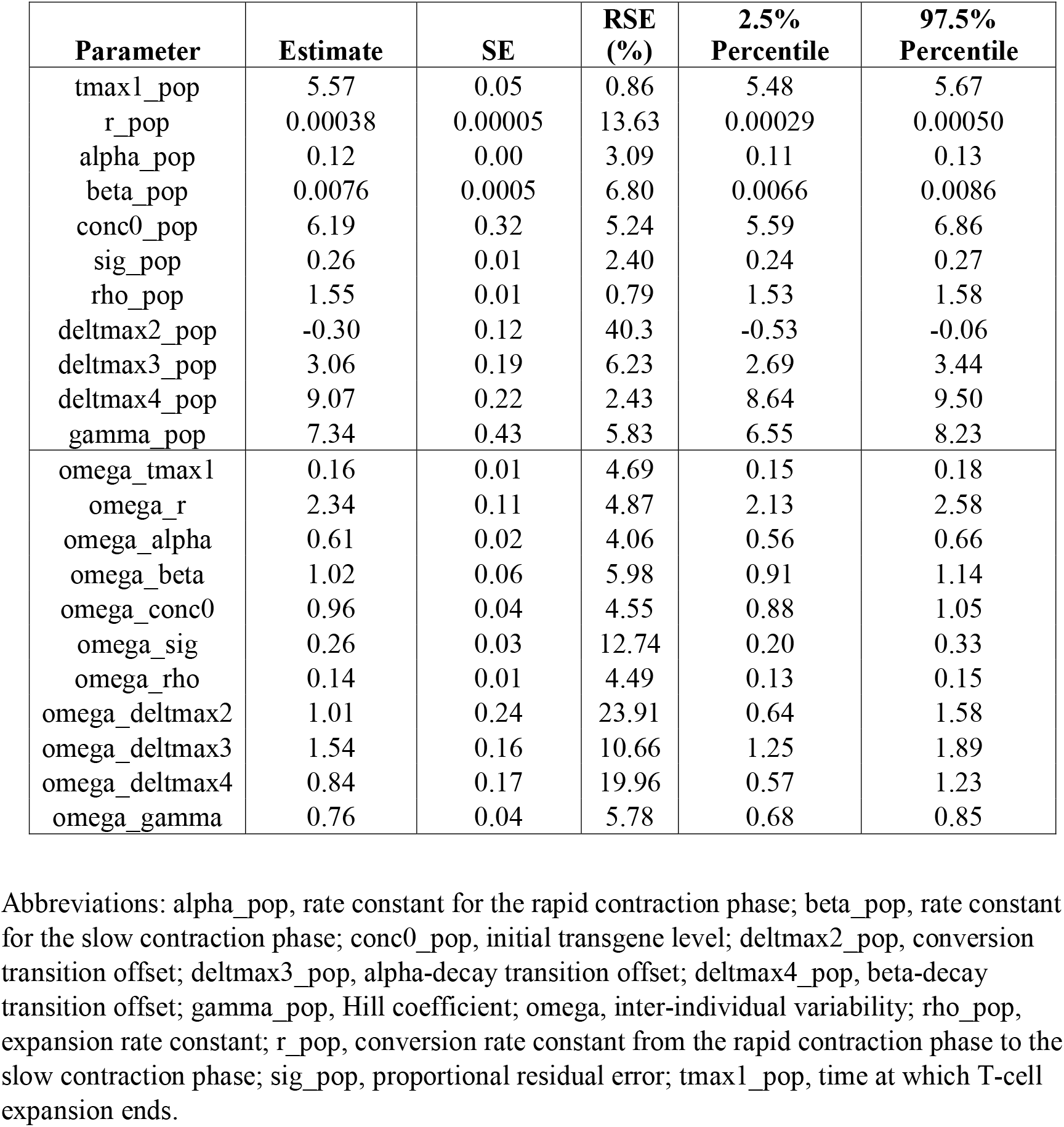
Summary of population cellular kinetic parameter estimates from the new model with smooth time-varying rate functions and process-specific transition timing.

### Full Bayesian confirmation (NONMEM Bayesian with Student’s t likelihood): robustness of inference for smooth gating and process-specific transition times

To obtain full posterior uncertainty for key parameters and timing offsets under a full Bayesian framework, we fit the process-specific transition-time model in NONMEM and examined posterior distributions (Figure 4). Figure 4 summarizes posterior distributions for kinetic parameters, residual dispersion, and timing parameters (tmax1 and deltmax2/3/4). Posterior summaries indicate that the conversion transition offset was credibly negative at deltmax2 = −0.36 days (95% credible interval (CrI): −0.51 to −0.23), supporting earlier conversion relative to the expansion transition. In parallel, the decay-related offsets were credibly positive, with tmax3 = 2.14 days (95% CrI: 1.68–2.48) and tmax4 = 7.23 days (95% CrI: 6.40–7.76).

**Figure 4.**
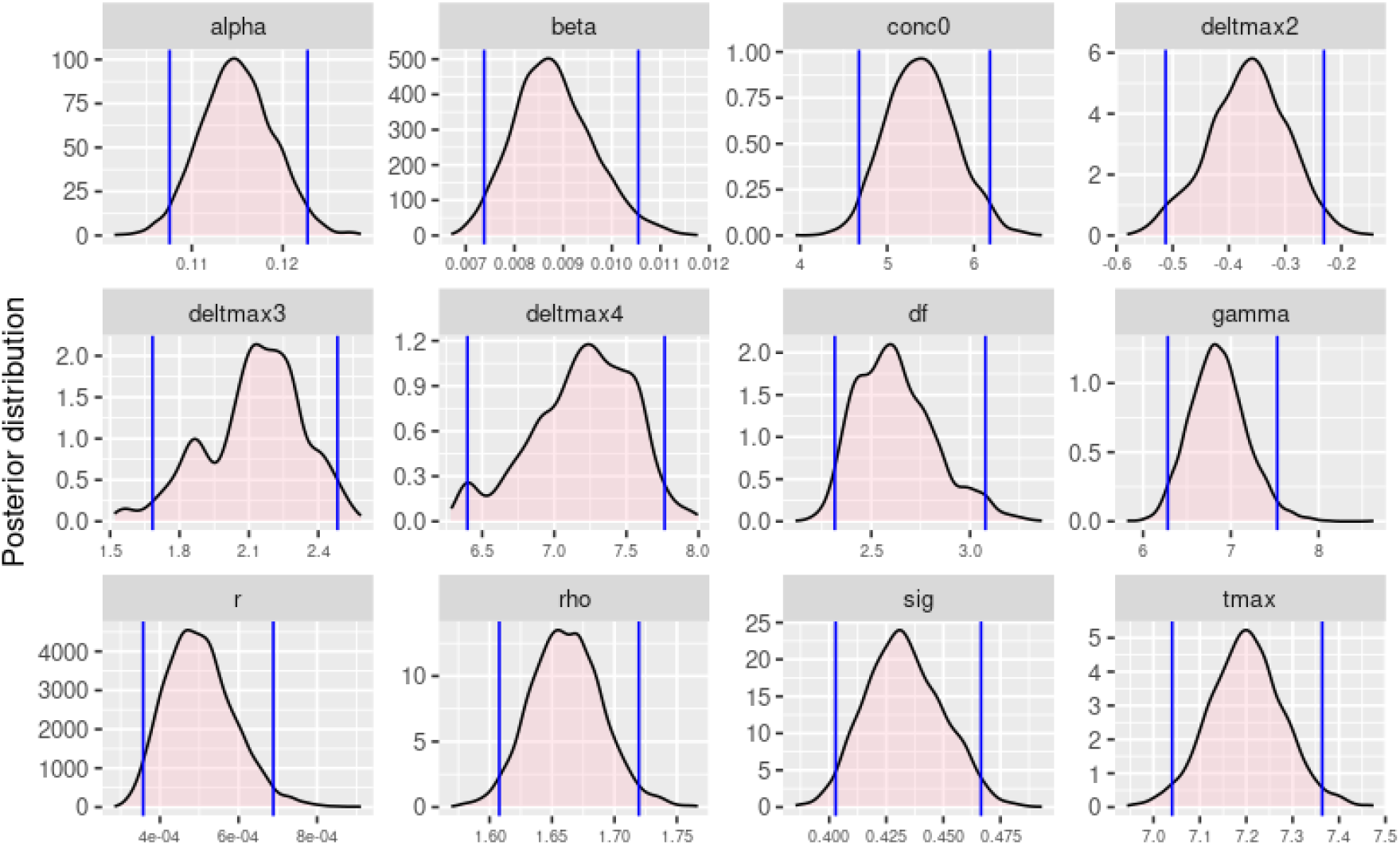
Posterior distributions of population parameter estimates from the full Bayesian analysis for the new model with smooth time-varying rate functions and process-specific transition timing. Blue vertical lines denote the 95% credible intervals of the posterior estimates. **alpha**, rate constant for the rapid contraction phase; **beta**, rate constant for the slow contraction phase; **conc0**, initial transgene level; **deltmax2**, conversion transition offset; **deltmax3**, alpha-decay transition offset; **deltmax4**, beta-decay transition offset; **gamma**, Hill coefficient; **omega**, inter-individual variability; **rho**, expansion rate constant; **r**, conversion rate constant from the rapid contraction phase to the slow contraction phase; **sig**, proportional residual error; **tmax1_pop**, time at which T-cell expansion ends; **df**, degrees of freedom.

Consistent with the Monolix output, the posterior distributions for the decay-related timing offsets (deltmax3 and deltmax4) were concentrated away from zero and aligned with the expected directionality.

Together, these posterior intervals provide coherent evidence that conversion initiates during expansion, whereas decay-related transitions occur later and are more strongly separated from the expansion transition.

## Discussion

CAR-T cell therapies represent a transformative class of adoptive immunotherapies with the capacity to expand, contract, and persist in vivo following a single infusion. ^4-6^ This kinetic behavior distinguishes CAR-T therapies from conventional small molecules and monoclonal antibodies and introduces modeling challenges that often require specialized statistical approaches. ^5, 7^ Relative to traditional therapeutics, CAR-T cellular kinetics profiles typically show larger inter-individual variability, more frequent influential observations, and a higher proportion of BLQ data, particularly during contraction and persistence. ^9-12, 15^ In this context, robust handling of both censoring and outliers is not a methodological nuance but a practical requirement for stable inference and credible prediction. ^13^

In this study, we addressed two limitations that frequently arise in CAR-T cellular kinetics modeling: (i) the practical burden of implementing robust residual error models—particularly when likelihood-based censoring contributions are required for BLQ observations—and (ii) the physiological oversimplification that can be introduced by forcing distinct biological processes to share a single synchronized transition time.

Our prior work highlighted the value of integrating Student’s *t*–distributed residuals with likelihood-based BLQ handling (M3 censoring) within a full Bayesian framework for CAR-T cellular kinetics modeling. ^9^ That combined approach was designed to address two common data features simultaneously: (1) BLQ observations that contain information about the lower tail of the distribution and should be included through a censoring likelihood rather than discarded or imputed, and (2) outlier contamination or influential observations that can destabilize estimation under Normal residual assumptions. Within that framework, three general conclusions emerged: M3 censoring enables principled use of BLQ information; heavy-tailed residuals improve robustness relative to Normal error models; and Bayesian methods offer coherent uncertainty quantification for complex nonlinear mixed-effects models.

A practical limitation of Student’s *t* likelihoods, however, is that their implementation— especially the CDF required for censored contributions—can be cumbersome **outside platforms such as NONMEM**, which provide built-in special-function support. This issue becomes immediately relevant when migrating a model across software ecosystems, such as translating a robust censored likelihood from NONMEM (where Student’s *t* and its CDF may be natively available) to Monolix (where user-implemented likelihood components may require explicit CDF evaluation). This work directly addressed that practical gap by evaluating the **Cauchy** distribution as an alternative heavy-tailed residual likelihood. Importantly, the Cauchy distribution is a specific case of the Student’s *t* distribution with one degree of freedom (ν = 1). ^24, 25^ The central question, therefore, was not whether Cauchy represents a fundamentally new robustness principle, but whether **Cauchy can serve as a functional surrogate** for Student’s *t* in settings where heavy-tailed robustness is needed and closed-form PDF/CDF expressions materially simplify implementation.

Across both simulation and real-data analyses, the results support that interpretation. In a controlled two-compartment IV PK simulation with terminal-phase outlier contamination, Cauchy residuals produced parameter recovery and individual-profile behavior that closely matched Student’s *t*, while avoiding the instability and bias observed under Normal residual assumptions as outlier severity increased. Importantly, Cauchy did not introduce meaningful distortion under clean data in this simulation setting, indicating that adopting a heavy-tailed residual likelihood did not incur an obvious penalty when the data were uncontaminated. In the integrated CAR-T case study under full Bayesian estimation with BLQ handled through a censoring likelihood, replacing Student’s *t* with Cauchy preserved posterior inference and yielded subject-level predictions that were largely indistinguishable from the Student’s *t* benchmark. From an implementation perspective, the closed-form CDF is operationally convenient when coding censored contributions, because BLQ likelihood terms require evaluation of probabilities such as P(Y ≤ LLOQ | model) (or two-sided limits, depending on the assay reporting convention). The availability of a closed-form CDF for Cauchy therefore provides a practical pathway for consistent robust + censored likelihood implementation across platforms.

Beyond the likelihood component, the second major focus of this work was structural: improving a widely used piecewise semi-mechanistic CAR-T model by (1) replacing discontinuous piecewise switching with **smooth S-shaped gating**, and (2) decoupling the assumption that multiple biological processes share a single synchronized transition time. Piecewise models are convenient and often effective, but they implicitly assume that biological processes switch instantaneously at a defined timepoint. In CAR-T biology, expansion, phenotypic evolution, contraction, and persistence are better represented as overlapping, gradual processes rather than synchronized on/off transitions. ^28, 29^ From a biological “continuum” perspective, cell differentiation is rarely instantaneous: the “switch” in a piecewise model implies that all cells change state at exactly *t*_max_, whereas in reality the population is heterogeneous—some cells transition early and others late. A sigmoidal (Hill-type) function provides a natural population-level representation of this gradual shift by allowing the effective rate to evolve smoothly over time. ^30, 31^ From a numerical-stability perspective, piecewise functions introduce discontinuities in derivatives, which can degrade the behavior of gradient-based estimation and sensitivity calculations, and can complicate convergence for commonly used algorithms. ^32, 33^ Smooth gating functions avoid these discontinuities, yielding differentiable transitions that better reflect gradual changes in net proliferation and loss while providing a flexible framework to represent heterogeneous and asynchronous cellular behaviors. Sigmoidal transitions are widely used to capture gradual phase changes in biological growth processes. ^27, 34^

A key observation from the process-specific transition model is that the three transition components are temporally decoupled. The decay-related offsets were consistently positive, indicating that decay-associated transitions occur later than the end of the cellular expansion. In contrast, the conversion-related offset was negative, consistent with the hypothesis that conversion from cytotoxic/effector-like to memory-like phenotypes begins **before** the expansion process fully terminates. Figure 5 shows that when transition timing is allowed to be process-specific, the inferred kinetics are not consistent with a single synchronized switch point. Instead, the expansion-rate term (rho) declines over a relatively narrow time window, whereas the conversion process (r) begins to rise earlier and the decay-related processes (*α, β*) activate later with broader transition windows. This pattern supports an asynchronous interpretation in which the observed CAR-T concentration–time profile reflects the superposition of partially overlapping biological programs—proliferation, differentiation, and contraction/persistence— rather than a single phase boundary shared by all processes. Biologically, the earlier onset of conversion (negative conversion-related offset; Figure 5) is consistent with evidence that effector–memory fate decisions can occur during the expansion phase, not only after peak expansion. CD8 T-cell differentiation is increasingly viewed as a continuum in which memory-precursor features can be established while cell division is ongoing, shaped by early antigen and cytokine signaling and supported by cellular heterogeneity generated during proliferation. ^35-37^ Accordingly, our model-based inference that a memory-like conversion program begins before proliferation fully terminates is physiologically plausible, and it is reflected by the smooth rise of *r* (*t*) prior to the complete shutdown of *rho* (*t*).

**Figure 5.**
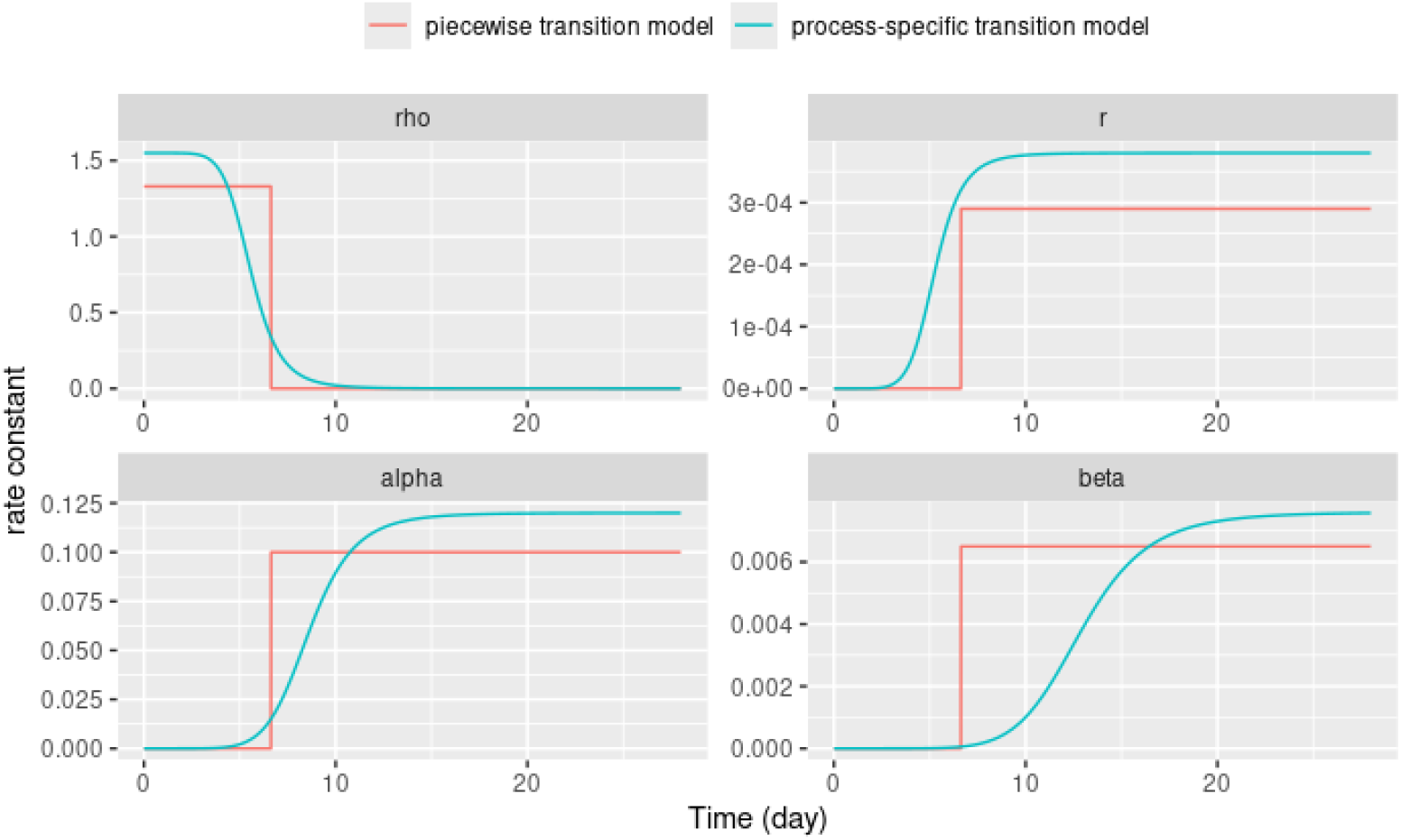
Comparison of piecewise versus process-specific transition timing for CAR-T kinetic rate constants (rho, r, alpha, beta) over time, based on population-typical rate functions computed from posterior means.

In contrast, the consistently positive offsets for *alpha* (*t*) and *beta* (*t*) align with the classical ordering of contraction and persistence, where cell-loss processes become dominant after the expansion peak and longer-lived subsets contribute to the slower terminal decline. ^7, 38, 39^ Beyond plausibility, this decoupling has practical implications: enforcing a single shared transition can structurally bias inference by misattributing early differentiation dynamics or delayed decay to the wrong phase, whereas process-specific timing provides a more interpretable bridge between estimated kinetic components and underlying biology and offers a clearer scaffold for future covariate or phenotype-informed extensions.

Two limitations merit emphasis. First, the Cauchy likelihood implies an extremely heavy-tailed residual distribution with fixed tail thickness, which may be more permissive than necessary for some datasets. A practical sensitivity analysis is therefore to compare Normal, Student’s *t* (with *v* estimated), and Cauchy models across key diagnostics and predictive checks to assess when Cauchy and low-*v* Student’s *t* yield similar versus meaningfully different inferences. Second, although smooth gating improves biological plausibility and may enhance numerical behavior, the additional transition parameters increase the need for careful identifiability assessment, particularly through a full Bayesian implementation with comprehensive posterior diagnostics. Future work could evaluate constrained parameterizations, informative priors, or joint modeling with relevant biomarkers to strengthen inference for transitions that are weakly informed by the available sampling schedule.

## Conclusion

This study advances CAR-T cellular kinetics modeling in two complementary ways. First, we show that a Cauchy residual likelihood can serve as a practical alternative for Student’s *t* in robust inference with BLQ censoring, offering comparable behavior with the implementation advantage of closed-form PDF/CDF for cross-platform portability. Second, we extend semi-mechanistic CAR-T models by replacing piecewise switching with smooth time-varying gating and by decoupling transition times across biological processes. The resulting framework supports an asynchronous interpretation of CAR-T kinetics in which conversion can initiate during expansion and decay-related transitions occur later, providing a more physiologically credible and flexible alternative to shared-Tmax piecewise architectures.

## Consent for publication

All the authors have reviewed and concurred with the manuscript.

## Funding

This work was sponsored and funded by Bristol Myers Squibb.

## Authors’ contributions

Y.C. and Y.L. contributed to conception and design; Y.C. and Y.L. contributed to acquisition of data; Y.C. and Y.L. contributed to analysis; all authors contributed to interpretation of data; Y.C. and Y.L. drafted and revised the article. Both authors made substantial contributions to conception and design, acquisition of data, or analysis and interpretation of data; took part in drafting the article or revising it critically for important intellectual content; agreed to submit to the current journal; gave final approval of the version to be published; and agreed to be accountable for all aspects of the work.

## Acknowledgments

The authors thank Robert J. Bauer from ICON for his assistance with the use of the NONMEM built-in density functions.

## Notes

### Competing Interest Statement

The authors have declared no competing interest.

